# Missing Value Imputation Approach for Mass Spectrometry-based Metabolomics Data

**DOI:** 10.1101/171967

**Authors:** Runmin Wei, Jingye Wang, Mingming Su, Erik Jia, Tianlu Chen, Yan Ni

## Abstract

**Introduction:** Missing values exist widely in mass-spectrometry (MS) based metabolomics data. Various methods have been applied for handling missing values, but the selection of methods can significantly affect following data analyses and interpretations. According to the definition, there are three types of missing values, missing completely at random (MCAR), missing at random (MAR), and missing not at random (MNAR).

**Objectives:** The aim of this study was to comprehensively compare common imputation methods for different types of missing values using two separate metabolomics data sets (977 and 198 serum samples respectively) to propose a strategy to deal with missing values in metabolomics studies.

**Methods:** Imputation methods included zero, half minimum (HM), mean, median, random forest (RF), singular value decomposition (SVD), k-nearest neighbors (kNN), and quantile regression imputation of left-censored data (QRILC). Normalized root mean squared error (NRMSE) and NRMSE-based sum of ranks (SOR) were applied to evaluate the imputation accuracy for MCAR/MAR and MNAR correspondingly. Principal component analysis (PCA)/partial least squares (PLS)-Procrustes sum of squared error were used to evaluate the overall sample distribution. Student’s *t*-test followed by Pearson correlation analysis was conducted to evaluate the effect of imputation on univariate statistical analysis.

**Results:** Our findings demonstrated that RF imputation performed the best for MCAR/MAR and QRILC was the favored one for MNAR.

**Conclusion:** Combining with “modified 80% rule”, we proposed a comprehensive strategy and developed a public-accessible web-tool for missing value imputation in metabolomics data.

## 1 Introduction

Metabolomics is the study of systematic identification and/or quantification of wide ranges of small molecule metabolites in bio-samples (cell, tissue, and biological fluids, etc.). Mass spectrometry (MS) is one of the main techniques for metabolomics studies (Dettmer et al. 2007). However, missing values, that certain compounds cannot be identified/quantified in certain samples, occur widely in MS-based metabolomics data due to technical and biological reasons (Bijlsma et al. 2006; Hrydziuszko and Viant 2012). Generally, there are three types of missing values, missing completely at random (MCAR), missing at random (MAR), and missing not at random (MNAR) (Gelman and Hill 2006; Little and Rubin 2002). Unexpected missing values are considered as MCAR if they originate from random errors and stochastic fluctuations during the data acquisition process (e.g., incomplete derivatization or ionization). MAR assumes the probability of a variable being missing depends on other observed variables (Gelman and Hill 2006; Little and Rubin 2002). Thus, missing values due to suboptimal data preprocessing, e.g., inaccurate peak detection and deconvolution of co-eluting compounds, can be called MAR. However, it is hard to distinguish these two types of missing values and some imputation methods can be applied for both MCAR and MAR data (Lazar et al. 2016). For targeted metabolomics studies, censored missing values caused by lower than the limits of quantification (LOQ) are considered as MNAR (Karpievitch et al. 2012). For example, previous study showed that bile acids exhibited large variations of concentrations in human serum, targeted identification of a panel of bile acids using MS technique produced many missing values due to LOQ (Xie et al. 2015).

The way of handling missing values in metabolomics data differs due to different sources. For those caused by random errors or stochastic fluctuations during the data acquisition process, the number of missing values is small and one can re-analyze or re-prepare bio-samples for data acquisition. For missing values produced during the data pre-processing step, filling peaks has been proposed in many tools by simply extracting and replacing with raw or baseline signals, e.g., fillPeaks in XCMS (Smith et al. 2006). However, these signals may not be accurate enough to represent real concentration levels of compounds unless baseline has been corrected and none co-eluting compounds exist nearby. For some missing values due to strict parameter settings during data processing, it is recommended to adjust or apply flexible parameter settings to retrieve real signals of certain compounds, such as low-concentration metabolites in biological samples (Y. Ni et al. 2016). Missing values due to LOQ are usually replaced with a small value or zero, which may lead to certain biases, e.g., distortions of the distribution of missing variables and underestimations of the standard deviation (Gelman and Hill 2006). Of course, in metabolomics, missing values that exist in more than 20% of samples may be removed directly from the data, which is called “80% rule” (Bijlsma et al. 2006). To decrease the risk of losing potential differential metabolites, “modified 80% rule” was proposed that variables are removed from the data when the proportion of non-missing elements are accounted for less than 80% among each group (Yang et al. 2015).

Instead of simply replacing missing with a specific value, more advanced imputation strategies have been proposed for handling missing values in –omics studies, such as k-nearest neighbors (kNN) imputation (Troyanskaya et al. 2001), random forest (RF) imputation (Stekhoven and Bühlmann 2012), and singular value decomposition (SVD) imputation (Hastie et al. 1999). Several software tools for metabolomics data analysis have implemented different methods dealing with missing values (Kessler et al. 2013; Luedemann et al. 2012; Xia et al. 2015; Katajamaa et al. 2006; Mak et al. 2014). MetaboAnalyst (Xia et al. 2009; Xia et al. 2015), one widely used metabolomics analysis toolkit, provides Probabilistic PCA (PPCA), Bayesian PCA (BPCA) and SVD imputation. However, the selection of methods for handling missing values can significantly affect subsequent data analyses and interpretations (Huan and Li 2015; Armitage et al. 2015), and it is difficult for users to decide an appropriate one for their data. Gromski et al. compared the performance of several missing value imputation methods on GC-MS metabolomics data, including zero, mean, median, kNN, and RF, and recommended RF as a favored one (Gromski et al. 2014). However, these imputation methods are suitable for MCAR/MAR only. Hrydziuszko et al. raised the importance of selecting optimal methods for treating missing values in metabolomics. They compared eight imputation methods in univariate and multivariate fashions and concluded kNN imputation was an optimal one (Hrydziuszko and Viant 2012). Although two types of missing values, MCAR/MAR and MNAR, were mentioned in their work, identical imputation strategies were applied and thus made it unclear to determine suitable methods for different types of missing values. The quantile regression imputation of left-censored data (QRILC), originally proposed for the imputation of MS-based proteomics data, imputes the left-censored missing in truncated fashion could be applied for MNAR in metabolomics (Lazar 2015). Thus, a comprehensive and systematic evaluation of different methods for handling missing values from different sources is needed for MS-based metabolomics studies.

In this study, we generated both MCAR/MAR and MNAR data in two separate clinical metabolomics studies, and then compared five different imputation methods (i.e., RF, kNN, SVD, Mean, Median) for MCAR/MAR and six imputation methods (i.e., QRILC, Half-minimum, Zero, RF, kNN, SVD) for MNAR. Then, we systematically measured the performance of those imputation methods using three different ways: (1) normalized root mean squared error (NRMSE) and NRMSE-based sum of ranks (SOR) were applied to evaluate the imputation accuracy for MCAR/MAR and MNAR correspondingly; (2) principal component analysis (PCA)/partial least squares (PLS)-Procrustes sum of squared error were used to evaluate the overall sample distribution; and (3) student’s *t*-test followed by Pearson correlation analysis was conducted to evaluate the effect of imputation on univariate statistical analysis. Results showed that RF imputation performed the best for MCAR/MAR and QRILC was favored one for MNAR. Finally, considering “modified 80% rule” together, we proposed a comprehensive strategy to deal with missing values in metabolomics studies with a public-accessible web-tool provided.

## 2 Material and Methods

### 2.1 Metabolomics data sets

Two real-world clinical metabolomics data sets were applied to evaluate the performance of different imputation methods. Since the measurements required comparisons between imputed data and original data, a complete raw data set was needed in our studies. Thus, we removed all missing values in our original data beforehand and left a complete data set for consequential analysis.

1. The first data set included a total of 977 de-identified subjects and 75 metabolites without missing values. It served as a large sample size data set for label-free evaluation.
2. The second data set was collected from a study of comparing metabolic profiles between obese subjects with diabetes mellitus and healthy controls (Yan Ni et al. 2015). After filtering all missing values, this data set contained a total number of 198 subjects (70 patients, 128 healthy controls) and 130 variables including metabolites (i.e., free fatty acids, amino acids, and bile acids) and/or derived ratios. It served as medium sample size data set for both label-free and labeled data evaluation.

### 2.2 Missing value generation

For MCAR/MAR generation, we randomly drew elements and replaced with missing values (NA) from the complete data matrix across the proportions from 1% to 50% in a step of 2.5% to generate a list of missing data sets.

For MNAR generation, according to our experience that the missing values usually occur in certain variables, we first randomly picked a certain number/proportion (from 4% to 80% in a step of 4%) of variables as missing variables. Then, we generated a random quantile cut off from the range 30%∼60% for each missing variable and replaced those elements under the cutoff with missing values. A list of MNAR data sets was then generated.

### 2.3 Missing value imputation methods

For the situation of MCAR/MAR, we applied five different imputation methods, which were:

- **kNN** (k Nearest Neighbors Imputation) (Troyanskaya et al. 2001): The original kNN imputation was developed for high-dimensional microarray gene expression data (n << p, n is the number of samples, and p is the number of variables). For each gene with missing values, this method finds the k nearest genes using Euclidean metric and imputes missing elements by averaging those non-missing elements of its neighbors. In metabolomics studies, we applied kNN to find k nearest samples instead and imputed the missing elements. We applied R package *impute* for this imputation approach.
- **RF** (Imputation with Random Forest) (Stekhoven and Bühlmann 2012): This imputation method applies random forest, a powerful machine learning algorithm, to build a prediction model by setting particular target variable with non-missing values as the outcome and other variables as predictors, then to predict the target variable with missing values iteratively. The R package *missForest* was used for this approach.
- **SVD** (Singular Value Decomposition Imputation) (Stacklies et al. 2007; Hastie et al. 1999): SVD imputation will initialize all missing elements with zero then estimate them as a linear combination of the k most significant eigen-variables iteratively until reaches certain convergence threshold. In metabolomics data, we scaled and centralized the data matrix first and then applied this imputation approach with the number of PCs setting to five by using R package *pcaMethods*.
- **Mean**: This method replaces missing elements with an average value of non-missing elements in corresponding variable.
- **Median**: This method replaces missing elements with a median value of non-missing elements in corresponding variable.

For the situation of MNAR, we applied RF, kNN, SVD and other three methods:

- **QRILC** (Quantile Regression Imputation of Left-Censored data) (Lazar 2015): QRILC imputation was specifically designed for left-censored data, data missing caused by lower than LOQ. This method imputes missing elements with randomly drawing from a truncated distribution estimated by a quantile regression. A beforehand log-transformation was conducted to improve the imputation accuracy. R package *imputeLCMD* was applied for this imputation approach.
- **Zero**: This method replaces all missing elements with zero.
- **HM** (Half of the Minimum): This method replaces missing elements with half of the minimum of non-missing elements in corresponding variable.

### 2.4 Performance evaluation

Normalized Root Mean Squared Error (NRMSE) has been commonly used to evaluate accuracy by calculating the differences between imputed values and real values by following formula (Oba et al. 2003)

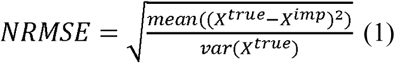

where X^*rue*^ is the true data, X^*imp*^ is the imputed data. We calculated NRMSE for the situation of MCAR/MAR on scaled data.

For MNAR, considering the missing value is not randomly distributed, using NRMSE directly might cause biased results especially for those imputation methods with determined values as we showed in the Supplements. Thus, we derived another metric, which was NRMSE-based sum of ranks (SOR). We first calculated the NRMSE for each missing variable and ranked them across different imputation methods. Consequently, we summed the rank of all missing variables for each method and made a comparison based on SOR. The SOR can be represented as following formula

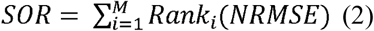

where M is the number of missing variables, *Rank_i_*(*NRMSE*) means the rank of NRMSE of different imputation methods in *i*_th_ missing variable. This non-parametric measurement provides a robust and unbiased comparison especially for the situation of MNAR.

NRMSE and SOR measured imputation accuracy in respect of the value of missing elements. Additionally, we measured the effects of different imputation methods in respect of overall sample distribution. To do this, we first applied the dimension reduction approach, e.g. PCA, to reduce the data dimensions. Then, Procrustes analysis, a statistical shape analysis, was conducted to compare the alteration of the imputed sample distributions with original sample distribution in the space of top PCs. Procrustes sum of squared errors were calculated as a quantitative measurements. R package *vegan* was applied for Procrustes analysis (Oksanen 2015).

In addition to the above label-free evaluations, we conducted extra evaluations from a statistical analysis perspective on the labeled data. These evaluations are mainly focused on the influences of different imputation methods on consequential statistical analyses. For univariate analysis, we conducted Student’s *t*-test on variables from imputed data and original data and *p*-values were then log-transformed considering their skewed distribution. We then conducted Pearson correlation analysis on the log *p*-values of imputed data and original data. For multivariate analysis, one of the most widely used methods in metabolomics studies is partial least squared regression (PLSR)/discriminant analysis (PLS-DA). In our case, since the phenotype outcome variable was case/control, we applied PLS-DA as a supervised dimensional reduction approach first. Then, we conducted Procrustes analysis to compare the sample distribution between imputed data and original data as we did previously. R package *ropls* was applied for PLS-DA (Thévenot et al. 2015).

## 3 Results

### 3.1 MCAR/MAR imputation and evaluation

We generated random missing values on both unlabeled and labeled data sets from the proportion of 2.5% to 50% in step of 2.5%. Five different imputation methods, RF, kNN, SVD, Mean and Median were conducted on all missing data sets. After imputations, NRSME were calculated by comparing differences between imputed data and complete data after z-score transformation (scaling and centralization). Z-score transformation makes it an unbiased comparison of using NRMSE considering the different ranges of variables in their original levels where the variable with large values will dominate the evaluation. Fig. 1a and b showed that RF imputation performed the best on both data sets with different proportions of missing values, followed by SVD and kNN imputation. We also found that kNN imputation method produced even larger NRMSE than two determined value imputation methods (i.e., mean and median) when the missing proportion increased to certain proportions. In contrast, these two determined value imputations performed stable on data with different proportions of missing values since the imputed “average” values made the mean squared error, the numerator of formula (1), equals/close to the denominator which is the variance of missing elements in complete data.

**Fig. 1.**
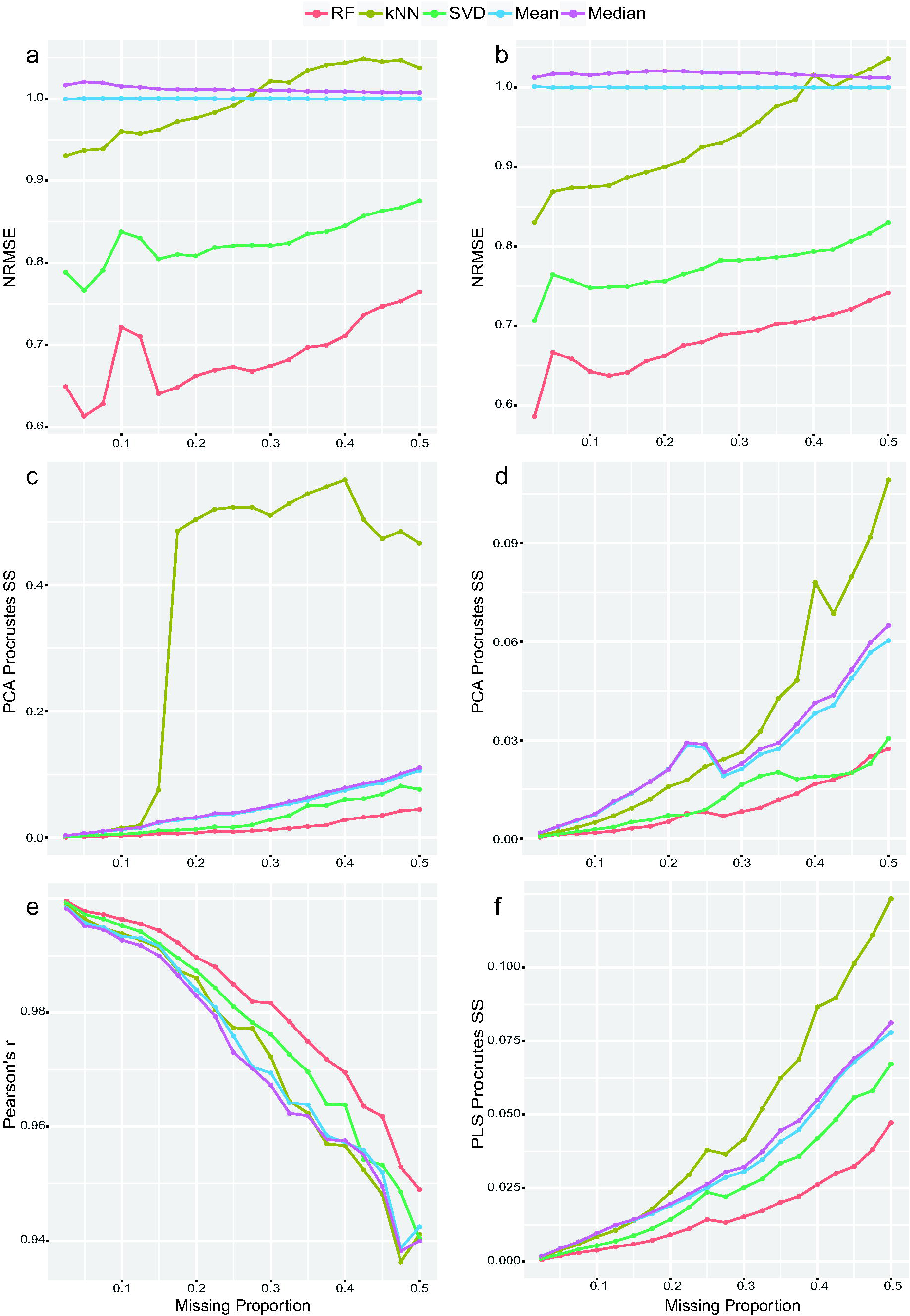
Evaluation of different imputation methods for MCAR/MAR. (**a-b**) NRMSE on unlabeled and labeled metabolomics data. (**c-d**) PCA-Procrustes sum of squared errors on unlabeled and labeled metabolomics data. (**e**) Pearson correlation of log *p*-values (*t*-test) of complete data and imputed data. (**f**) PLS-Procrustes sum of squared errors.

Next, we applied PCA to both complete and imputed data sets and selected first two PCs as they represented the most variance. Then, we applied Procrustes analysis to compare the distribution of sample points on top two PCs of imputed data with the complete data using scaled sum of squared errors. Less distortion of imputed data (smaller sum of squared errors) represented a better recovery of imputation regarding the original sample distribution. Results showed that RF performed the best across different missing proportions on both data sets (Fig. 1c and d). Meanwhile, these two determined value imputation methods, mean and median, gave very similar trends. In contrast, kNN started performing the worst once the proportion of missing values reached to certain cutoff values.

With sample phenotype information (case/control), we performed both Student’s *t*-test and PLS-DA on the labeled data set with or without missing values. First, we conducted *t*-test on each variable. Then, we measured the Pearson’s *r* between the log *p*-values calculated from imputed data and complete data. Results showed that RF imputation maintained the highest correlation coefficients across different missing proportions (Fig. 1e), which indicated that the most information of original univariate results had been remained using RF method. In addition, we applied PLS-Procrustes analysis and found that RF imputation kept the best among five imputation methods with the lowest sum of squared errors (Fig. 1f).

### 3.2 MNAR imputation and evaluation

For each MNAR data set, six different imputation methods were applied with two aims: first, to evaluate the performances of those imputation methods previously applied on MCAR/MAR (i.e., RF, kNN, SVD) on the situation of MNAR; second, to compare the performance of three left-censored imputation methods (i.e., QRILC, HM and Zero) on MNAR. After imputation, we applied SOR to evaluate their imputation accuracy performances. Results (Fig. 2a and b) showed that all three imputation methods, RF, SVD, and kNN, performed poorly on MNAR, together with Zero imputation that had been commonly used in metabolomics data analysis. In comparison, QRILC produced much smaller SOR values followed by HM imputation, showing consistent good performances on data with different numbers of missing variables.

**Fig. 2.**
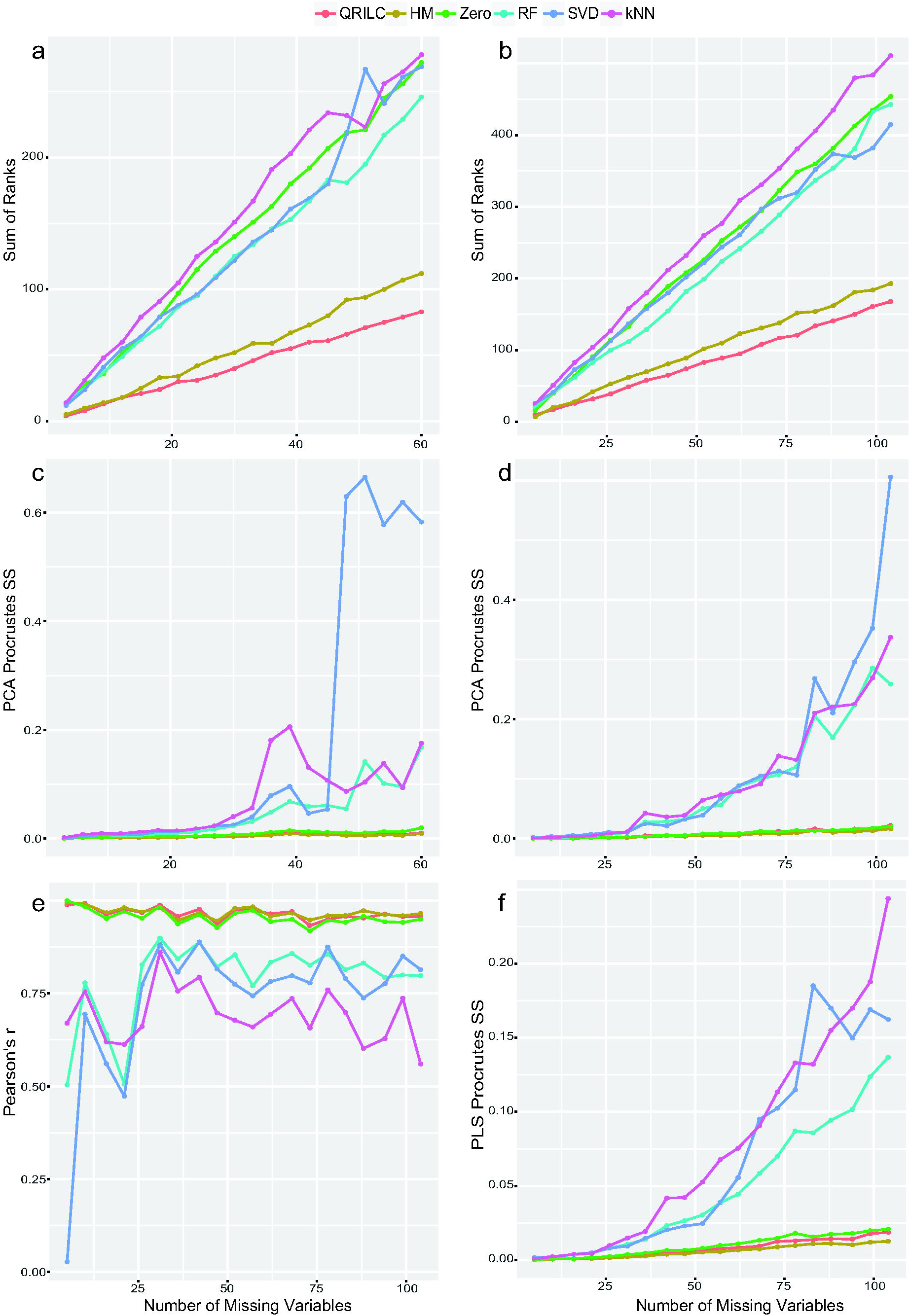
Evaluation of different imputation methods for MNAR. (**a-b**) SOR on unlabeled and labeled metabolomics data. (**c-d**) PCA-Procrustes sum of squared errors on unlabeled and labeled metabolomics data. (**e**) Pearson correlation of log *p*-values (*t*-test) of missing variables from complete data and imputed data. (**f**) PLS-Procrustes sum of squared errors.

From PCA-Procrustes analysis, we observed that three imputation methods for MCAR/MAR changed original sample distribution to a large extent as the number of missing variables increased (Fig. 2c and d). In contrast, three MNAR imputation methods showed very consistent results, among which, HM performed slightly better across the different number of missing variables followed by QRILC.

For the correlation analysis on log *p*-values of missing variables, Fig. 2e also demonstrated that three MNAR imputation methods performed better with higher correlation coefficients and QRILC performed almost as good as HM. For the PLS-Procrustes analysis (Fig. 2f), the result was similar to PCA-Procrustes analysis that we observed from Fig. 2d. To summarize, both QRILC and HM showed decent and stable performances across different numbers of missing variables.

### 3.3 QRILC and HM imputation

Next, we further compared the overall performance of HM and QRILC methods. QRILC imputes the left-censored data by randomly drawing values from a truncated normal distribution while HM replaces missing elements by using the half of the minimum of non-missing values. As a determined value imputation method, HM has limitations in some circumstances (e.g., distorting distributions and underestimating the variances of missing variables that further affect multivariate analysis) (Gelman and Hill 2006). In this work, we randomly selected ten variables from the unlabeled data set to construct a new data set and assigned eight of them as missing variables which contained 40% ∼ 80% left-censored missing values. PCA analysis on complete data, QRILC imputed, and HM imputed data with top 2 PCs showed that QRILC kept the overall shape and distribution of original data set while HM showed a subgroup of samples gathering around to a straight line (Fig. 3a-c). This was because missing values of those samples were replaced by the same determined values, making those samples gathered towards a straight line on the PCA score plot. In addition, the violin plots also demonstrated that HM method severely distorted the distributions of eight variables (Fig. 3d-k).

**Fig. 3.**
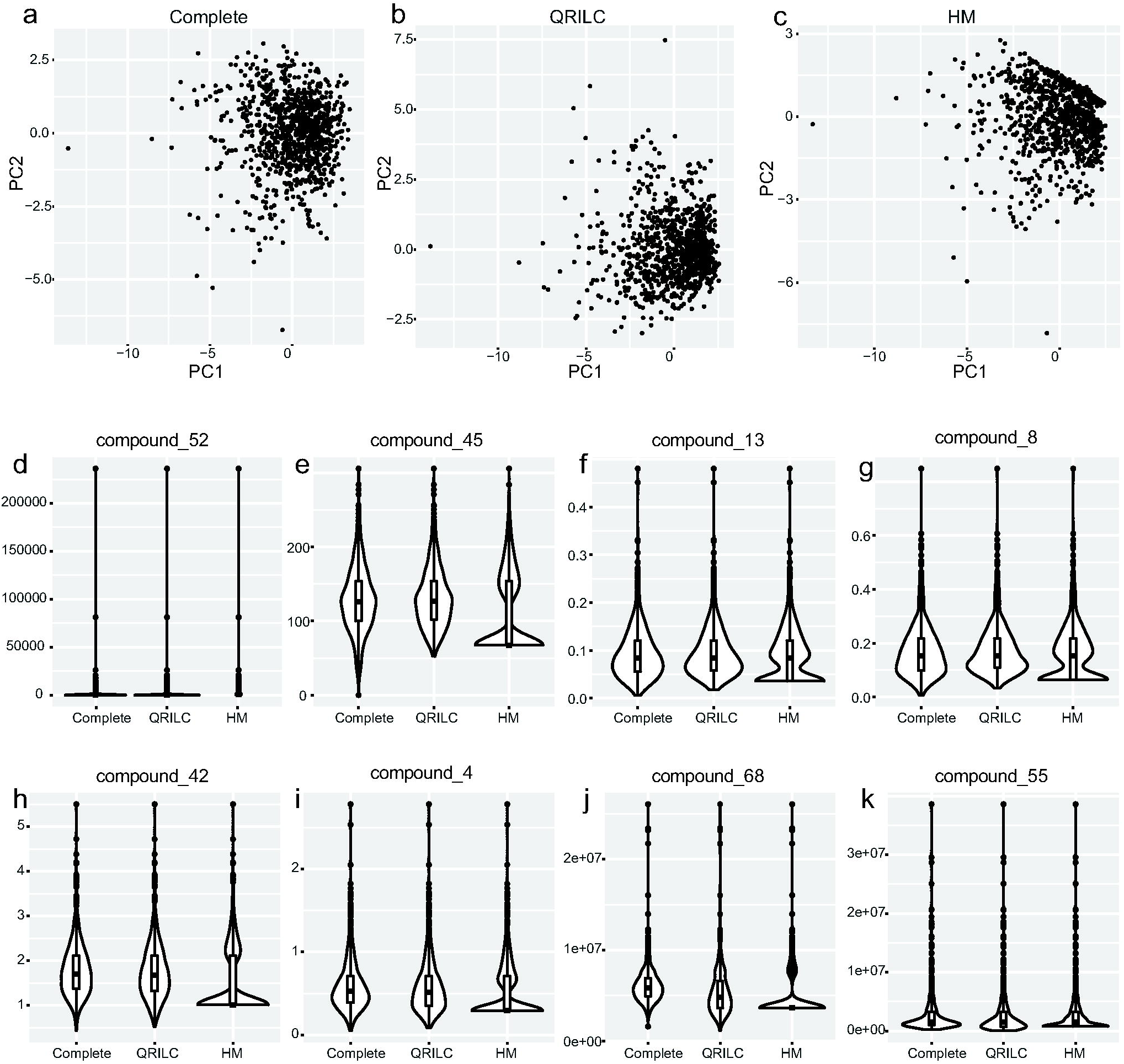
Comparisons between QRILC and HM for MNAR. (**a-c**) PCA score plot for complete data, QRILC imputed data and HM imputed data on top 2 PCs. (**d-k**) Violin plots of eight missing variables.

## 4 Discussion

To avoid potentially biased comparisons, we have explored the performance of each imputation method for MS-based metabolomics data and optimized parameter settings or data pre-processing steps to reach optimal performance. For example, kNN was previously applied on a p × n gene expression matrix (genes in rows and samples in columns) while we found kNN performed better on an n × p metabolomics data matrix (samples in rows and metabolites in columns). The underlying difference is that instead of finding k nearest variables, which are genes in their original case, we tried to find k nearest samples to represent the missing ones. This is reasonable since gene expression data usually contains a large number (more than 10,000) of genes where co-expression occurs frequently thus neighbored genes are usually good representation for missing ones. For SVD imputation, as suggested by the original paper, a pre-scaling of the data will increase the accuracy, and we also found using top five PCs is a good choice in our study rather than the default setting of two. Since the original scale function in R is irreversible, we implement a scale-recover function, which enables us to recover the scaled table to the original scale. For QRILC imputation, log-transformation beforehand was found useful not only to improve imputation accuracy but also to ensure a positive value in the original scale.

In this work, we systematically evaluated a total of eight imputation methods on different types of missing values in terms of imputation accuracy, sample distribution, and statistical analysis. Results shown that RF imputation performed the best for MCAR/MAR data. For MNAR, we found that QRILC and HM performed better than others, however, HM could distort the distribution of single variables as indicated by violin plots or of a linear combination of variables as indicated by PCA score plots. Thus, we recommend the more smoothed method QRILC for MNAR due to LOQ, especially in targeted metabolomics studies. Combing classic “modified 80% rule” together, we proposed a comprehensive strategy to deal with missing values in metabolomics studies (Fig. 4). For both targeted and profiling/non-targeted metabolomics data, we recommend (1) checking raw data and if necessary, adjusting parameter settings in order to fill back certain missing values in an accurate way; (2) applying “modified 80% rule” to remove those unreliable variables that have more than 20% missing values in each subgroup of samples; (3) for the remaining missing values, users need to decide the possible reasons and choose an appropriate one for missing value imputation; (4) for targeted metabolomics data, QRILC imputation is recommended for missing values due to LOQ and for the profiling/non-targeted metabolomics data, RF imputation is the recommended method to be applied for MCAR/MAR.

**Fig. 4.**
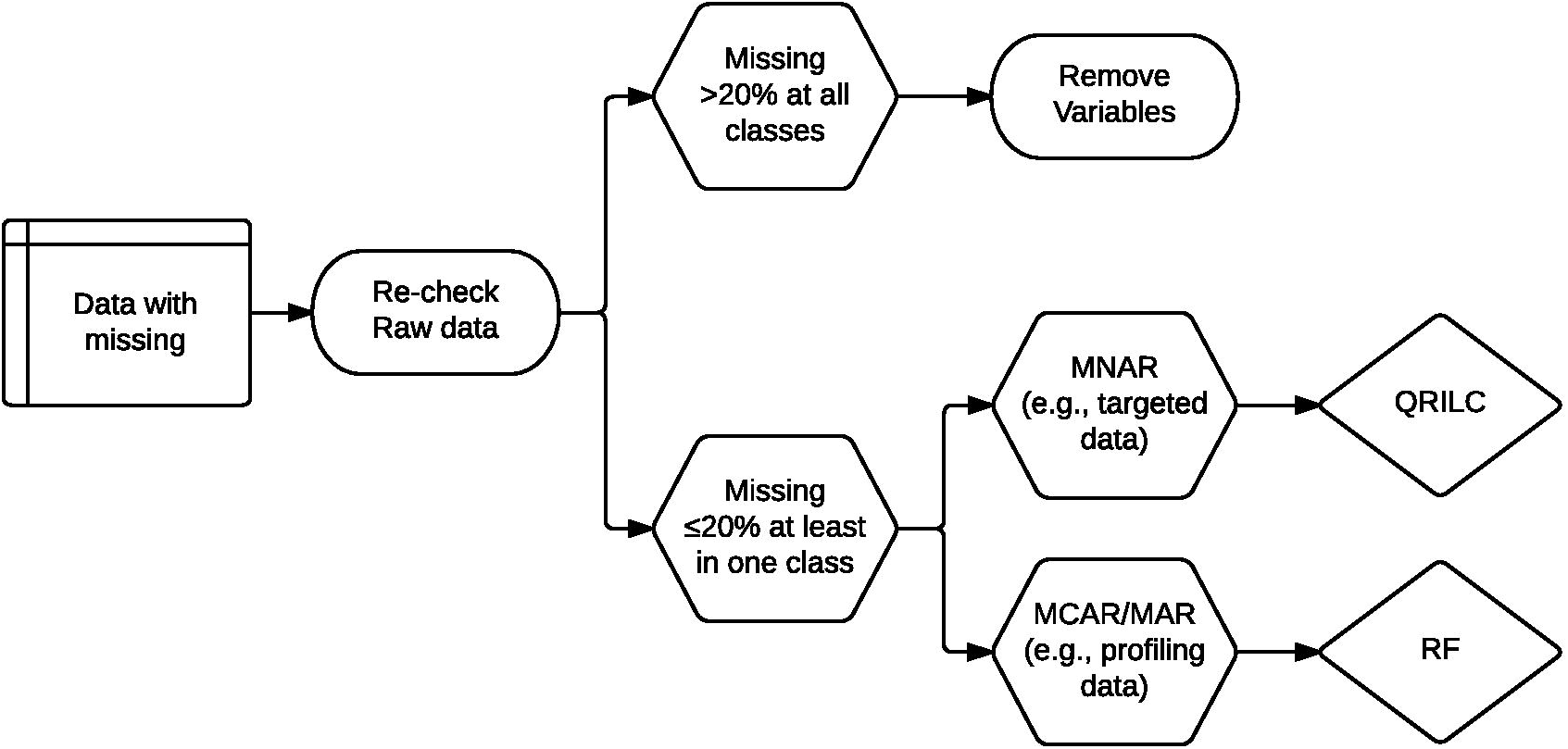
An imputation strategy for metabolomics studies.

In addition, a web-tool (https://metabolomics.cc.hawaii.edu/software/MetImp/) has been developed allowing users to upload their data and choose an appropriate method for missing value imputation. As described in this work, eight different methods dealing with missing values are provided in the web server. “Modified 80% rule” was the default setting for group-wise missing filtering while RF and QRILC were provided as default imputation methods corresponding to MCAR/MAR and MNAR. Additionally, all settings are interactively modifiable for the flexibility of usage. Finally, a complete table with all missing values being imputed will be generated and users can download for further statistical analyses.

There are various imputation methods in the field and new methods are developing by researchers sequentially. To assist future researches in metabolomics data imputation, we packaged our evaluation processes (including missing generation and imputation, evaluation with NRMSE, SOR, correlation, PCA/PLS-Procrustes analysis and visualization) into R functions and developed a convenient and comprehensive imputation evaluation pipeline (https://github.com/WandeRum/MVI-evaluation). This evaluation pipeline enable researchers to perform further and specific studies on missing value imputation problems, e.g., comparing new imputation methods with existing ones, and evaluating different methods on specific data sets. In the future, new methods dealing with metabolomics missing values, especially for MNAR, will be introduced for comprehensive comparison through our evaluation pipeline and added to our web-tool.

## 5 Conclusion

Missing values occur widely in MS-based metabolomics data due to technical and biological reasons. Three different types of missing values, MCAR, MAR, and MNAR, are commonly occurred in metabolomics studies. In this work, using two separate metabolomics data sets, we systematically evaluated eight different imputation methods, in terms of imputation accuracy, sample distribution, and statistical analysis. We found that RF imputation performed the best for MCAR/MAR data and QRILC was favored one for MNAR imputation. Combing “modified 80% rule” together, we proposed a comprehensive strategy to deal with missing values in metabolomics studies. Finally, a public-accessible web server has been developed for dealing with metabolomics missing value imputation.

## Associated Content

Supplements.docx – Imputation Evaluation Vignette

## Acknowledgements

We acknowledge valuable contributions, input, and discussions from Professor Wei Jia.

